# Exploring Protein Patterns, Cavity Interactions, and Therapeutic Insights in Cancer

**DOI:** 10.1101/2025.06.03.657615

**Authors:** Paloma Tejera-Nevado, Belén Otero-Carrasco, Alejandro Rodríguez-González

## Abstract

Protein sequence alignments are essential for identifying proteins’ shared structural and functional features. Detecting short amino acid sequences, termed patterns, across lung cancer and other related datasets facilitates the identification of relevant features. This study builds on previous findings by exploring proteins that share common patterns already identified. Using sequence matching at 5% and 10% occurrence thresholds, we identified 2,368 and 47 patterns, respectively. To reduce complexity and refine the dataset, shorter patterns from the 10% occurrence streamlined the analysis by isolating highly relevant patterns while reducing redundancy among proteins sharing sequence segments. Subsequent analyses integrated structural predictions for protein folding comparison, enabling the detection of patterns in different proteins and the identification of potential key residues. During cavity detection prediction, some amino acids were inspected in detail to assess their impact on protein function and their relevance in drug-target interactions. These insights were considered during docking studies, focusing on proteins used in treatments with pre-described ligands. By connecting raw sequence data to folding structures and functional features, we identified critical protein cavities that underscore the role of mutations in altering protein behavior and influencing drug-target interactions. These findings highlight protein activity’s structural foundations and their importance in understanding cancer biology. By uncovering conserved sequence patterns and their structural implications, this study provides insights into potential biomarkers and therapeutic targets, that could aid in developing more effective cancer treatments.

## I. INTRODUCTION

Proteins, made of 20 amino acids, are crucial for structural and enzymatic functions across four structural levels [1, 2]. Understanding their sequences is key to disease research, helping identify patterns, mutations, and therapeutic targets. Protein-ligand docking refers to a computational technique used to predict how a small molecule (the ligand) binds to a protein (the receptor or target) [3]. This process is crucial for drug discovery and understanding molecular interactions.

Lung cancer, the leading cause of cancer-related deaths, is mainly caused by smoking, with most cases diagnosed late. Early detection improves survival, and the WHO stresses prevention through tobacco control and reducing environmental risks. It includes Non-Small Cell Lung Cancer (NSCLC), a slower-growing type, and Small Cell Lung Cancer (SCLC), a more aggressive form, with 1.8 million deaths recorded in 2022. Other risk factors are second-hand smoke, pollution, and genetic predisposition [4]. NSCLC is a group of lung cancers comprising at least three distinct histological types: squamous cell carcinoma, adenocarcinoma, and large cell carcinoma, all classified as non-small cell lung cancers. These cancers typically respond poorly to conventional chemotherapy [5].

Cancer develops when genetic mutations accumulate in key genes, particularly those that regulate cell growth, division (proliferation), and the repair of damaged DNA. Somatic mutations in the TP53, EGFR, and KRAS genes are frequently observed in lung cancer. Additionally, mutations in several other genes are commonly related to cell proliferation, overseeing the maturation of cells to perform specific functions (differentiation) and regulating cell death [6]. NR4A3 is important in lung and breast cancer because it acts as a tumor suppressor, both dependent and independent of p53. It is directly activated by p53, which enhances its expression. NR4A3 suppresses cancer cell proliferation and promotes apoptosis by upregulating pro-apoptotic genes. Additionally, NR4A3 interacts with the anti-apoptotic protein Bcl-2, preventing it from blocking apoptosis. High NR4A3 expression is linked to better survival outcomes in cancer patients, highlighting its potential as a key factor in cancer progression and therapy [7]. NR4A3 has been identified as an oncogenic driver in acinic cell carcinoma (AciCC) of the salivary glands due to a recurrent genomic rearrangement. Its immunostaining has been reported to have diagnostic value, particularly in high-grade and challenging cases [8].

Patterns help identify key sequence features, such as binding and active sites, by analyzing conserved regions in multiple sequence alignments. In the present study, patterns were defined as short amino acid sequences identified across different proteins through sequence alignment. These sequences contained at least four amino acids, considering this the minimum relevant length [9]. The patterns identified in the target protein sequences associated with treatments for NSCLC, which served as the foundation for this research, were derived from a previous study [10]. In that study, an in-house computational methodology was developed to identify shared amino acid sequences across different protein sequences, followed by docking analysis to uncover significant relationships in depth. Relevant patterns in reference and target proteins were identified through shared and non-overlapping analysis. Specifically, this approach aims to assess the relevance of these patterns in the quaternary structure of proteins and their relationship with drug-binding sites, comparing proteins from other cancer types, including breast, colon, pancreas, and head and neck cancers. The paper is organized in the following order: Section 2 describes the tools and resources used to study protein patterns. In Section 3, the results obtained are presented and discussed in Section 4. Finally, Section 5 summarizes the conclusions of this study and highlights potential avenues for future research.

## II. MATERIAL AND METHODS

### A. Descriptive Analysis of Patterns in Lung Cancer

An initial descriptive analysis of the data under consideration was conducted. A previous study identified relevant patterns in NSCLC protein targets at two different occurrence thresholds: 10% and 5 % [10]. These limits correspond to the minimum frequency with which patterns had to appear in the target proteins of drugs for this disease.

To begin, 5%-occurrence patterns were filtered using 10%-occurrence patterns, considering both identical and similar patterns. The number of patterns shared between the target proteins of lung cancer drugs and those of other studied cancer types was calculated. The patterns were quantified across different datasets to identify similarities between the various cancer types (breast, colon, pancreas, and head and neck cancers). To illustrate the overlap of these patterns, a Venn diagram was employed. Additionally, an UpSet plot was utilized to further analyze the distribution and overlapping of patterns, providing a detailed view of how these patterns are shared and distributed across the datasets.

### B. Protein Sequences and Structural Predictions

DISNET is a database that integrates information from various sources on biological, phenotypic, and pharmacological data [11]. DISNET has been previously used for works in the area of disease understanding [12] and drug repurposing for rare diseases [13], COVID-19 [14], and others [15-18]. The protein IDs correspond to the UniProt entry identifiers [19]. To obtain the predicted protein structures, two sources were used: structures deposited in the AlphaFold DB [20, 21] and the predictions generated using the AlphaFold server (https://alphafoldserver.com/), which provides five different output models. Information about the action types of the ligands was retrieved from DrugBank portal (https://go.drugbank.com/).

### C. Protein-Ligand Docking Analysis Tools

CB-Dock2 is an advanced protein-ligand blind docking tool that integrates cavity detection and homologous template fitting [22]. In CB-Dock2, the CurPocket ID refers to a unique identifier for specific cavities or binding pockets within a protein structure. These pockets are identified as potential sites for ligand binding, and the CurPocket ID helps guide the docking process by focusing on relevant regions of the protein for ligand analysis. The Vina score reflects the binding affinity of a ligand to a protein, with more negative values indicating stronger binding. It is generated using the AutoDock Vina algorithm during blind docking, where the protein’s binding pockets are explored without prior knowledge of the binding site.

COACH-D enhances protein-ligand binding site prediction by combining the consensus algorithm of COACH with molecular docking via AutoDock Vina, offering refined ligand poses with reduced steric clashes [23]. The Score in COACH-D is a confidence score that ranges from 0 to 1, indicating the reliability of a predicted ligand-binding site, with higher scores suggesting more accurate predictions.

The docking prediction outputs, including the protein-ligand complexes in PDB format, were visualized and analyzed using ChimeraX (v. 1.6.1) [24]. The Matchmaker tool [25] was used to compare treatment proteins with candidate proteins, allowing the visualization of pattern locations. Additionally, the hydrophobicity feature enables surface visualization, with atoms color-coded based on their hydrophobic properties. This facilitated the analysis of cavities and ligand positioning.

## III. RESULTS

### A. Descriptive Analysis of Patterns in Lung Cancer

At the 5% occurrence threshold, 2,368 relevant patterns were identified as common between the target proteins of lung cancer drugs and those of the other studied cancer types. These patterns were longer and present in a smaller subset of proteins. However, due to the low occurrence threshold, the resulting list was too extensive for comprehensive evaluation. Conversely, at the 10% occurrence threshold, 47 relevant patterns were identified, shared among a larger set of proteins. Based on these results, the first step in this study was to filter the 5%-occurrence patterns using the 10%-occurrence patterns, considering not only identical patterns but also similar ones. A similar pattern is defined as one that contains the core pattern but may include additional amino acids at either end, forming a longer pattern. This filtering process resulted in a new subset of 37 relevant patterns, which comprised longer patterns potentially providing greater biological insights and were associated with a reduced subset of proteins. Among these 37 patterns, one identical pattern, “DTLS”, and 35 similar patterns were identified.

Subsequently, the number of patterns shared among the different cancer types was analyzed to identify the most shared patterns. The results were visualized using a Venn diagram (Fig. 1), which displays both unique patterns associated with each cancer type and those shared across multiple types. Unique patterns included eight in breast cancer, one in colon cancer, and none in pancreas or head and neck cancers. In contrast, nine patterns were found to be shared among all four cancer types: “AAVAG”, “AEEEE”, “DTLS”, “EEAEK”, “EEEED”, “KEKEK”, “PPSPP”, “PSPPP”, and “RLQAL”.

**Fig. 1.**
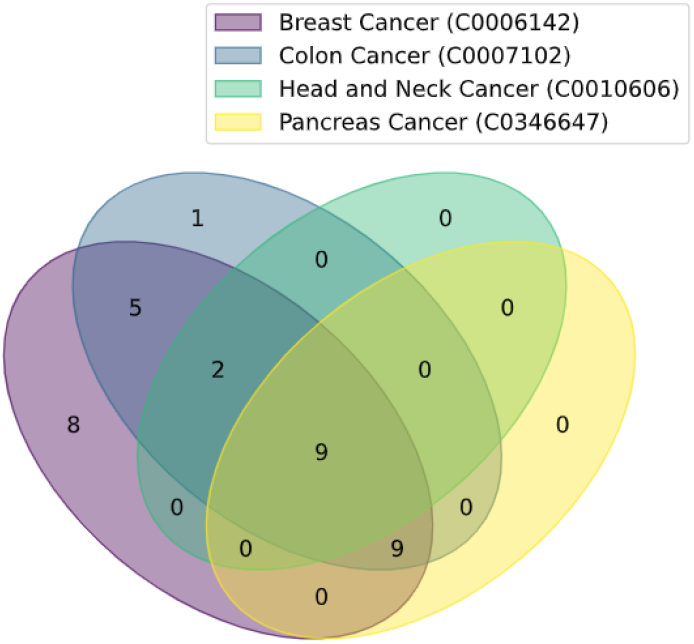
Venn diagram showing the overlap of patterns in Breast, Colon, Pancreas, and Head and Neck cancers. Terms in parentheses indicate a Concept Unique Identifier (CUI) from the Unified Medical Language System^®^ (UMLS^®^) Methathesaurus. Each CUI represents a specific concept, linking synonymous terms across different source vocabularies. CUIs are formatted as the letter ‘C’ followed by seven digits.

Additionally, Table 1 outlines all possible combinations of shared patterns across the considered cancer types. As the number of combined cancers increases, the detected patterns decrease, with the four-cancer combination having the lowest count, as well as the combinations involving pancreas and head and neck with breast and colon.

**TABLE I.**
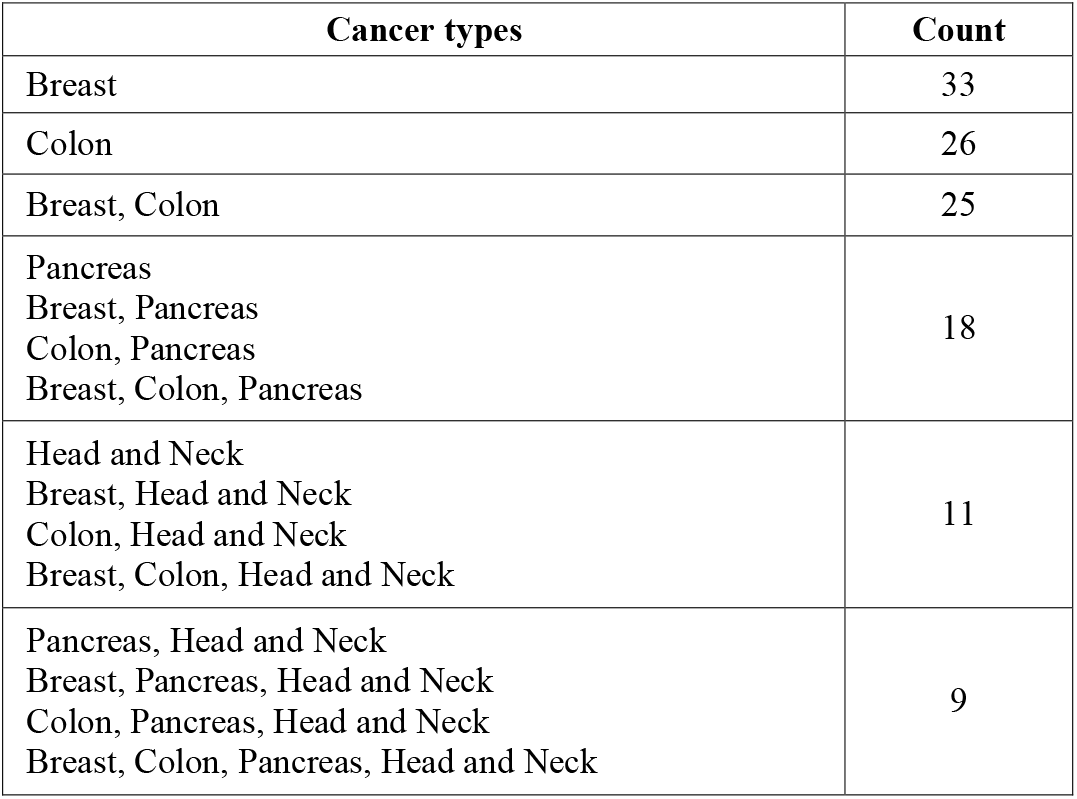
Number of detected patterns in the cancer types.

An analysis was also conducted to investigate patterns of interest associated with each target protein in NSCLC treatments and determine whether different target proteins shared the same pattern. This descriptive analysis was visualized using an UpSet plot (Fig. 2), which reveals that the “DTLS” pattern was shared among five different target proteins, representing the maximum value observed. It can be seen in this same graph that of the 37 selected patterns, 26 of them are shared between at least two target proteins.

**Fig. 2.**
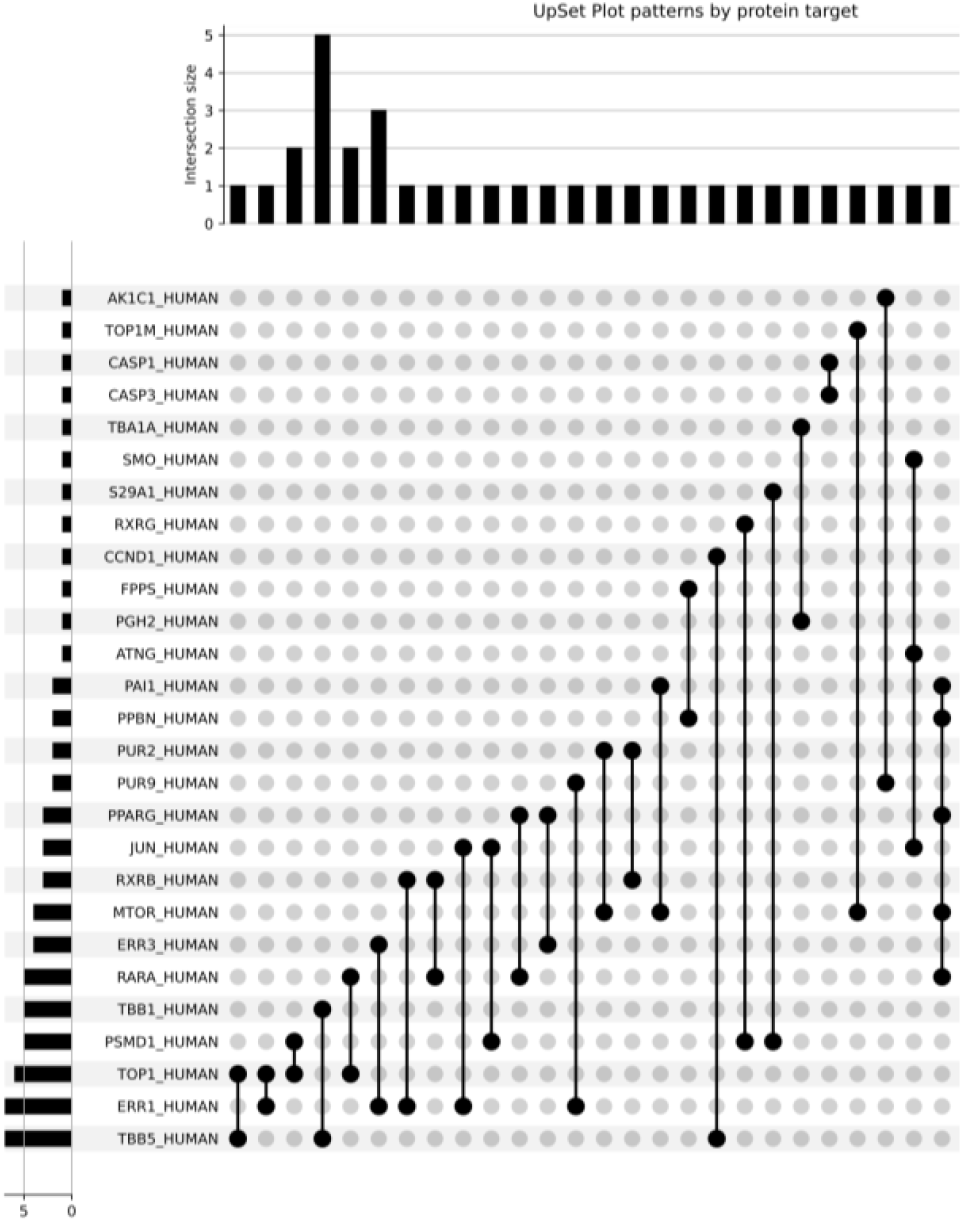
UpSet plot illustrating the distribution and overlap of patterns across various protein targets, highlighting the shared patterns between them.

### B. Analysis of shared patterns between proteins

In a further study of possible amino acid coincidences between proteins, the protein of interest NR4A3 (Q92570) was included in the search. The hypothesis focuses on whether longer and shared patterns can be detected between proteins. By examining the detected pattern “PSPP” in the 10% occurrence and using the NR4A3 protein, three longer patterns were identified among the 5% occurrence candidates: “PPSPP”, “PSPPP”, and “VPSPP”.

“PPSPP” is present in the proteins RARA (P10276) and RXRB (P28702) from the treatment dataset and is associated with the drugs isotretinoin and bexarotene, respectively. A total of 35 proteins share this pattern between the proteins from the treatment dataset and those from the other four cancer types. This analysis included NR4A3 and NR4A2 (P43354), and the pattern corresponds to an unstructured region. The protein RXRB and the drug bexarotene were used to search for cavities using CB-Dock2 and COACH-D, and it was observed that the pattern was not included in the detected cavities (starting amino acid positions: 75 for RARA and 97 for RXRB, respectively). However, the output data showed a sequence of alanines in a row, starting from amino acid position 46 of RXRB (Fig. 3). The “AAAAA” pattern was also detected in NR4A3 in the obtained results. Further inspection of this protein revealed that the pattern is not part of a cavity.

**Fig. 3.**
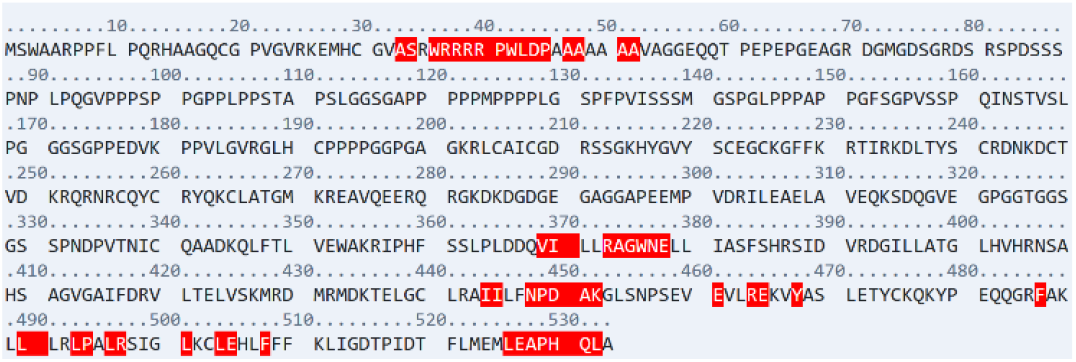
Sequence and amino acids involved in cavity detection for RXRB. Structure-based cavity detection with CB-Dock2 identified pockets in the input structure, submitted as the protein RXRB (AF-P28702). Amino acids involved in CurPocket C1 were identified as A47, A48, A51 and A52, and were marked in red.

Initially, only one protein folding prediction from AlphaFold DB was included. However, a detailed analysis was later conducted using all five outputs provided by the AlphaFold server, which could reveal small differences in the spatial distribution of amino acids. The “PPSPP” pattern was not detected as part of a cavity in the NR4A3 and NR4A2 proteins (data not shown). In contrast, the “AAAAA” pattern was identified as part of a cavity in CurPocket C1 of RXRB, derived from the treatment dataset. However, it was not detected in NR4A3, which belongs to proteins from other cancer types that share this pattern. The NR4A2 sequence does not contain this pattern.

A structural analysis of RXRB using CurPocket identified five distinct cavities (C1-C5) with varying volumes, sizes, and binding affinities (Table II). Cavity C1 was the largest, with a volume of 4459 Å^3^, while C5 was the smallest at 254 Å^3^. The Vina scores, indicating binding affinity, ranged from –12.2 for C2 (strongest binding) to –5.9 for C5 (weakest binding). The cavities also varied in their spatial properties, with different center coordinates and dimensions, influencing their potential role in ligand interactions. The five CurPockets were selected for blind docking. C1, which contained the alanine pattern, had a Vina score of –7.7, ranking second. However, none of the alanine residues were present. The following amino acids were identified: I370, A374, A451, K452, G453, R497, L501, L504, E505, F508, A528, H530, Q531, L532.

**TABLE 2.** Bexarotene - RXRB (AlphaFoldDB) - CB-Dock2.

| CurPocket ID | Cavity volume (Å<sup>3</sup>) | Center (x, y, z) | Cavity size (x, y, z) | Vina score | Docking size (x, y, z) |
| --- | --- | --- | --- | --- | --- |
| C1 | 4459 | -3.3,-9 | 21,30,19 | -7.7 | 22,35,22 |
| C2 | 1036 | -14,6,13 | 16,19,23 | -12.2 | 22,22,28 |
| C3 | 387 | -4,4,21 | 11,10,6 | -6.8 | 22,22,22 |
| C4 | 313 | 4,9,8 | 10,12,6 | -7.6 | 22,22,22 |
| C5 | 254 | -18,-14,0 | 7,10,8 | -5.9 | 22,22,22 |

To assess the consistency of pattern detection in cavity identification, five outputs of the RXRB protein structure prediction were obtained from the AlphaFold server. Cavity detection in model 3 suggested that A47 and A48 could be potential residues involved in the “AAAAA” pattern. Docking simulations were performed on the five pockets (Table III). In the case of pocket C1, the contact residues included W42 L43 D44 P45 A48 V311 K314 SER315 Q317 G318 D344 K345 Q346 F348 T349 A398 T399 Q531. The A48 is detected in this sequence.

**TABLE 3.** Bexarotene - RXRB (model3) - CB-Dock2.

| CurPocket ID | Cavity volume (Å<sup>3</sup>) | Center (x, y, z) | Cavity size (x, y, z) | Vina score | Docking size (x, y, z) |
| --- | --- | --- | --- | --- | --- |
| C1 | 4894 | -7,11,8 | 20,29,30 | -9.2 | 22,34,35 |
| C2 | 538 | 4,4,21 | 11,12,15 | -7.8 | 22,22,22 |
| C3 | 340 | 7,3,0 | 7,13,12 | -8.0 | 22,22,22 |
| C4 | 327 | 21,-36,14 | 8,14,7 | -5.9 | 22,22,22 |
| C5 | 162 | 3,18,-1 | 8,7,6 | 3.2 | 22,22,22 |

The five CurPocket cavities identified in RXRB varied in volume, size, and binding affinity. Cavity C1 had the largest volume (4894 Å^3^) and a Vina score of –9.2, making it the most favorable. C2 and C3 had smaller volumes (538 and 340 Å^3^, respectively), with Vina scores of –7.8 and –8.0. Cavity C4 was slightly smaller (327 Å^3^) and had a lower Vina score of – 5.9, while C5 was the smallest (162 Å^3^) with a much weaker Vina score of 3.2. Docking sizes for all cavities were consistent, except for C1, which had a slightly larger docking size.

### C. Identifying longer patterns through noverlapping analysis

Another approach used in pattern detection was to identify longer patterns by combining them with a non-overlapping approach.

In this analysis, RARA (P10276), a protein included in the treatment dataset, was used as a reference for further investigation. A search was conducted within the 5% occurrence datasets to detect overlapping patterns. The patterns “TLLK” and “LLKA” were identified at positions 259 and 260, respectively, sharing the subsequence “LLK”. The intersection of these patterns led to the selection of 12 candidate proteins from the other dataset. The amino acid sequence was predicted to form an alpha-helix structure.

Two types of analyses were performed: first, the detection of cavities followed by a comparison with closely related proteins, such as RARG (P13631) and RARB (P10826); second, a comparison with NR1D1 (P20393). The protein structure predictions included are from AlphaFold DB.

The proteins RARA, RARB, RARG, and NR1D1 were docked with isotretinoin, and the analysis identified overlapping patterns of amino acids in their cavities. The method used CB-Dock 2, helped determine the binding sites and Vina scores, indicating the strength of these interactions (Table IV). The patterns observed for RARA, RARB, and RARG showed similarities, especially with amino acids like T259, K262, A263, L266, D267, and E415, which were common across these proteins. The Vina scores suggested that the binding affinity was comparable among these three, with RARB showing the strongest interaction, followed by RARA and RARG. NR1D1 shows the highest score of −7.4, and one amino acid from the pattern was identified: K473.

**TABLE 4.** Docking analysis of protein-ligand interaction using CB-Dock 2: Vina scores and amino acid involvement.

| Protein | CurPocket ID | Vina Score | Amino acids involved |
| --- | --- | --- | --- |
| RARA | C2 | -6.9 | **T259 L261 K262 A263** A264 C265 L266 D267 R339 Q340 D341 R385 S388 A389 A392 M413 L414 E415 N416 S417 E418 |
| RARB | C3 | -7.0 | **T259 K262 A263** L266 D267 R339 Q340 D341 L384 R385 S388 A389 A392 E415 N416 S417 E418 |
| RARG | C1 | -6.6 | I260 **T261 K264 A265** L268 D269 R341 M342 D343 R387 S390 T391 A394 M415 N418 P419 E420 |
| NR1D1 | C2 | -7.4 | G12 V13 I14 T15 Y16 I17 F443 T444 V447 R448 **K473** H602 L606 |

In the context of protein-ligand docking simulations using COACH-D method, the analysis focused on detecting binding patterns and comparing CScore values across different proteins (Table V). The results show that RARA and RARB share a similar binding pattern, with CScore values of 0.72 and 0.76, respectively, involving amino acids like L261 and K262. RARG (CScore: 0.68) presents a distinct pattern, with amino acids shared in different positions. NR1D1, with the lowest CScore of 0.17, shows a unique set of amino acids, including L472 and K473, as part of the analyzed pattern.

**TABLE 5.** Docking analysis of protein-ligand interaction using COACH-D: Vina scores and amino acid involvement.

| Protein | Rank | CScore | Amino acids involved |
| --- | --- | --- | --- |
| RARA | 1 | 0.72 | V240 K244 F249 I254 Q257 I258 **L261 K262** P408 L409 E412 M413 |
| RARB | 1 | 0.76 | V240 K244 F249 I254 Q257 I258 **L261 K262** P408 L409 E412 M413 |
| RARG | 1 | 0.68 | V242 K246 F251 I256 Q259 I260 **L263 K264** P410 L411 E414 M415 |
| NR1D1 | 1 | 0.17 | R448 V451 K455 Q465 H466 Q468 V469 **L472 K473** |

The predicted protein folding structure of NR1D1 was visualized with hydrophobic labels (Fig. 4). The pattern “TLLKA”, corresponding to an alpha-helix, includes the amino acids L472 and K473, which are part of this pattern and were detected in the docking simulations at selected pockets.

**Fig. 4.**
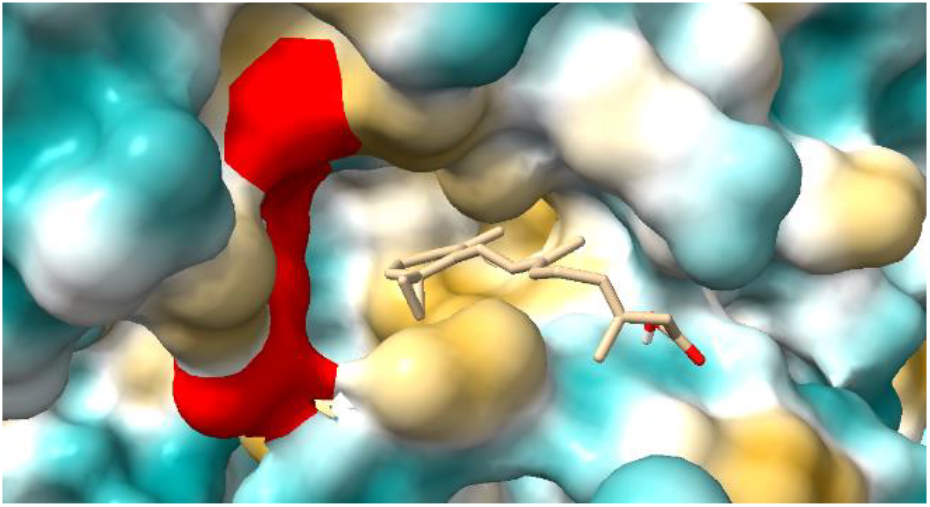
The predicted protein folding structure of NR1D1, visualized with hydrophobic labels. The pattern “TLLKA” corresponds to an alpha-helix, and the amino acids L472 and K473, belonging to the pattern and detected in the docking at the selected pockets, are colored red. A detailed view shows the predicted cavity and isotretinoin docking in NR1D1.

## IV. DISCUSSION

This study presents a novel integration of docking analysis with amino acid patterns identified from protein comparisons in lung cancer, linking them to treatment insights and proteins from four distinct cancer types. The first nine patterns “AAVAG”, “AEEEE”, “DTLS”, “EEAEK”, “EEEED”, “KEKEK”, “PPSPP”, “PSPPP” and “RLQAL”, were found to be shared among all four cancer types. “PPSPP”, in particular, was used as the starting point for the study, analysis, and evaluation of cavity detection (Fig. 1). Additionally, the number of patterns detected in each cancer type differs: 33 for breast cancer, 26 for colon cancer, 18 for pancreas, and 11 for head and neck cancer (Table I). The fact that the cancer types seem to maintain the same order as the counts for the patterns detected and the number of proteins in each dataset (before removing those already present in the lung cancer treatment dataset) could suggest that more patterns are found when more proteins were already included from the beginning. Specifically, the dataset includes 3,062 proteins related to breast cancer, 1,905 related to colon cancer, 728 related to pancreatic cancer, and 121 related to head and neck cancer.

To identify potential candidates for further analysis, the proteins sharing some patterns were determined. PAI1, PPBN, PPARG, MTOR, and RARA shared the “DTLS” pattern, present in their sequence (Fig. 2). An initial inspection was conducted for the proteins corresponding to the “DTLS” pattern. However, due to the large number of candidates, a more thorough screening is necessary, as the first inspection made it challenging to pinpoint the most relevant protein candidates for cavity detection and docking analysis. Notably, the cancer dataset, which includes four different cancer types, contained 48 proteins with this pattern, while 5 distinct proteins from the lung cancer treatment dataset should be further analyzed with their ligands, adding to the complexity of the required analysis.

Another important factor to consider is that RARA also shares patterns with RXRB, and their respective drugs, isotretinoin and bexarotene, have been identified. A study suggests that isotretinoin may interact with Fox1, which could help explain some of its currently unexplained effects [26]. The pharmacological action is not fully understood. Moreover, isotretinoin acts as an agonist of RARA, but additional mechanisms of action remain unclear. In addition, bexarotene acts as an agonist of RXRB. It specifically functions as a DNA-binding transcription activator, RNA polymerase II-specific. Considering this, it was relevant to explore cavity detection and possibilities for docking.

By using different docking tools, some of the amino acids included in the patterns within the protein sequences were detected as part of cavities and docking sites close to the ligand. Tools like CB-Dock2 offer improved accuracy and efficiency for drug discovery and bioinformatics research [22]. Building on template-based approaches, FitDock [27] employs hierarchical multi-feature ligand alignment to improve protein-ligand docking accuracy and speed, complementing tools like CB-Dock2 by focusing on docking efficiency and alignment optimization. COACH-D [23] complements template-based like CB-Dock2 and FitDock. In the first scenario, where the “PPSPP” pattern is found at positions 75 and 96 for RARA and RXRB, respectively, it was not detected as part of the cavities or involved in ligand interactions in any of the five protein prediction structures determined by AlphaFold (data not shown). This pattern was also identified in the candidate proteins NR4A3 and NR4A2, at positions 391 and 357, respectively. The pattern forms part of an unstructured region of the protein and does not form part of the cavity of amino acids involved in the predicted docking sites. It has been described that methionine, alanine, leucine, glutamate, and lysine exhibit a strong tendency to form α-helix structures, whereas proline and glycine have low helix-forming propensities [28]. Moreover, the “AAAAA” pattern was detected, starting at positions 46, 47, or 48, and it was also identified by the common pattern identification method. Notably, this pattern is part of an alpha helix. In contrast, NR4A3 contains this pattern at position 227, where it is unstructured, and it is not present in NR4A2. Cavity detection results identified some of the alanines implicated in the cavity (Fig. 3), but this was observed only for RXRB when bexarotene was included as a ligand.

The data used in this study include proteins, their sequences, and the diseases with which they are associated. NR4A3 has been linked to breast cancer, while NR4A2 is associated with colon cancer. NR4A3 and NR4A2 are the primary driver genes of acinic cell carcinoma (AciCC). Additionally, the simultaneous overexpression of NR4A3 and MYB characterizes a subset of AciCC patients with high-grade transformation, which is associated with an exceptional prognosis [29]. The expression of NR4A3 in breast acinic cell carcinoma and its similarity to salivary AciCC have been studied [30]. Breast and salivary AciCC are distinct with different molecular drivers, unlike other salivary gland-like tumors found in the breast. Some genes are involved in different types of diseases, and the underlying mechanism may vary. Therefore, analyzing different proteins that share certain patterns could help identify similarities and variations in their structures. NR4A2 is a gene essential for dopaminergic neuron differentiation, and mutations contribute to dopaminergic dysfunction in Parkinson’s disease [31]. Taking everything into account, it will be necessary to further study predicted cavities and determine if there are amino acids detected in other patterns, particularly those associated with these proteins.

It is necessary to estimate metrics, as the comparison involves not only different tools but also various predictions (Tables II–III). While RMSD (root mean square deviation) assesses deviations using a reference protein, the variability of proteins in this study allows only the localization of the ligand, cavity, and pattern amino acids in the target protein (Tables IV–V). The final part of the analysis includes proteins that contain the pattern “TLLKA”. Specifically, RARB, RARG, and NR1D1 are associated with breast cancer, while RARB is also linked to colon cancer. After inspecting the protein RARA, which is involved in treatment with isotretinoin, it was detected that NR1D1 shares some amino acids and was detected in the docking at the selected pockets (Fig. 4), opening new insights for this type of in silico analysis.

## V. CONCLUSION AND FUTURE WORK

By comparing protein sequences from previously described lung cancer treatment targets with those associated with other cancer types, we identified amino acid sequences that are shared between them, referred to as patterns. Comparing potential docking sites from the original ligand and the target proteins allowed us to assess whether the amino acids in these patterns were also part of the cavities, or if they provided insights into how closely related the proteins are. Different prediction outputs and docking tools were used for comparison. The information generated suggests that these findings can provide valuable insights and enhance our understanding, particularly before validation in experimental setups. This work is ongoing, intending to develop a workflow that can be used to infer information about three-dimensional structures and surface interactions. It will also aim to determine whether ligands or drugs interact directly with proteins or form part of complexes and variants, features critical in biotechnology, including drug repurposing.

## Acknowledgment

The work is a result of the project “Data-driven drug repositioning applying graph neural networks (3DR-GNN)” that is being developed under grant “PID2021-122659OB-I00” from the Spanish Ministry of Science and Innovation.

